# Comparison of Permanent and Resorbable Alginate Microspheres for Safety and Embolic Effects in a Porcine Renal Artery Embolization Model

**DOI:** 10.1101/2025.01.15.633170

**Authors:** Sankalp Agarwal, Masum Pandey, Ronan Duggan, Con O’ Brien, Andrew. L. Lewis, Brendan Duffy, Joan McCabe, Liam Farrissey

**Affiliations:** Centre for Research in Engineering and Surface Technology, TU Dublin, Ireland; CrannMed Ltd, Galway, Ireland

## Abstract

**Purpose:** This study evaluated porcine renal artery embolization using resorbable alginate microspheres versus permanent microspheres (Embozene), assessing vessel recanalization, local/systemic foreign body reactions, and potential complications.

**Materials and Methods:** Twelve Yorkshire swine (3 female, 9 male) underwent unilateral kidney arterial embolization with either Embozene or resorbable microspheres. Angiography was performed to confirm embolization. Vessel recanalization for was determined through angiography. The effect of embolization was evaluated by assessment of serum chemistry markers and by renal histopathology at 1, 3, 7 and 28 days post-procedure.

**Results:** Embolization procedures with both embolic agents were performed with 100% technical success. Hematology, coagulation factors and serum chemistry were largely unremarkable throughout the study period of 28 days. Alginate resorbable alginate microspheres facilitated compete recanalization and revascularisation within the initial 24hr observation window, while the permanent microsphere remained present at all evaluated time points. No inflammation was observed with resorbable microsphere whereas Embozene-induced inflammation was minimal to mild. Vascular histopathologic alteration with resorbable microspheres was limited and resolved by 24 hours, whereas Embozene caused sustained injury extending alteration to the internal elastic lamina, tunica media or adventitia.

**Conclusion:** Alginate resorbable microsphere demonstrated effective temporary embolization with a recanalization time of around 24 hours together with complete renal tissue recovery, no systemic impact, and minimal local reaction compared to Embozene. The findings suggest that embolization using resorbable microspheres is safe and may be suitable for use where temporary embolization of vessels is required.

## 1 INTRODUCTION

In musculoskeletal (MSK) embolization for pain relief, clinicians have found that transient occlusion alone can be therapeutically effective[1]. However, no embolic agents are currently FDA-cleared for this specific use in the U.S., leading to the off-label use of materials like Gelfoam and imipenem/cilastatin (IMP/CS) antibiotics[2], [3]. These agents exhibit inconsistent degradation profiles, influenced by factors such as embolization extent,vascular anatomy and the inflammation status of the target site [4], [5] [6]. This highlights the need for a purpose-built, resorbable embolic agent with predictable performance.

Alginate-based microspheres have long been explored for embolization, both alone and as drug carriers [7] [8]. These are prepared by ionotropic gelation using divalent ions like Ca^2^□, but they are highly stable in vivo and unsuitable for temporary embolization. However, studies have shown that alginate microspheres can be degraded on demand using calcium chelators or alginate lyase [9] [10].

The current study adopted an ingenious strategy wherein freeze-dried alginate lyase enzyme loaded calcium-crosslinked alginate microspheres were prepared [11]. Upon rehydration and administration into the body, the dormant enzyme is reactivated, enabling controlled self-degradation of crosslinked-alginate matrix[11]. To evaluate the potential as a temporary embolic agent in humans, the biological safety of resorbable microspheres (RM) must first be assessed, which is ordinarily achieved by comparison with a previously characterized predicate product. Herein we report on the safety and embolization effects over a 28-day period of RM in comparison with permanent embolic microspheres (PM, Embozene®) in a porcine renal artery embolization model.

## 2 Materials and Methods

### 2.1 Preparation of sterile freeze-dried RM (SakuraBead®)

In brief, 2% w/v sodium alginate solution in 0.1□M pH□2.8 acetate buffer was mixed with 0.15U alginate lyase to achieve the desired degradation time. Using an electrostatic encapsulator(Nisco, Switzerland), the mixture was dispensed through a 0.09□mm stainless steel ID needle into a 2% w/v calcium chloride bath at 3□mL/h under 3□kV for 30 minutes to optimize the production of ∼200□µm beads. Beads were washed with cryoprotectants [12], freeze-dried at −45□°C for 90 hours, vacuum-sealed, and sterilized using gamma radiation (> 10 kGy) to achieve Sterility Assurance Level (SAL) of 10^−6^ [11].

### 2.2 Animal Model and Husbandry

Twelve healthy Yorkshire swine (n=3 females, n=9 males) were used under FDA Good Laboratory Practices (GLP; 21 CFR Part 58) with IACUC approval (Project No. I00362). Animals were housed with daily access to Purina Lab Diet (#5084) and water ad libitum.

### 2.3 Study Design

The porcine renal embolization model was chosen for this safety study, as it has been previously used in similar studies. [13] [14].This study was conducted as per GLP guidance at CBSET, USA, with a protocol agreed with FDA focused on overall safety to minimise the use of animals (3R’s), rather than powered to deliver statistical significance in observed effects. Each pig underwent a single secondary branch renal artery embolization procedure on Day 0, with either RM or PM (Embozene®, 100 µm, (Varian Medical Systems, USA), as outlined in Table 1.

**Table 1:** Study design outline.

| Treatment Group | Number of Animals | Description of Treatment | Number of Treatments per Animal | Necropsy Timepoint |
| --- | --- | --- | --- | --- |
| 1 | 2 | Kidney injection with RM | 2 | 24 Hours |
|  | 1 | Kidney injection with PM |  |  |
| 2 | 2 | Kidney injection with RM | 2 | Day 3 |
|  | 1 | Kidney injection with PM |  |  |
| 3 | 2 | Kidney injection with RM | 2 | Day 7 |
|  | 1 | Kidney injection with PM |  |  |
| 4 | 2 | Kidney injection with RM | 2 | Day 28 ± 1 |
|  | 1 | Kidney injection with PM |  |  |

As the primary motivation for the development of the RM was for MSK indications, 100 µm Embozene was selected as the appropriate comparator product because it has been used in human trials in the US for the same indications (ClinicalTrials.gov ID: NCT03491397), Fewer PM interventions were performed compared to RM, as the embolic effects of the former are well-established in the literature [15]. Histopathology and haematology analyses were performed pre- and post-embolization at 24 hours, Day 3, 7 and 28.

### 2.4 Renal embolization using PM and RM

For the PM group, 100□µm Embozene® microspheres (Figure 1 D) were prepared as per the manufacturer’s Instruction For Use (IFU), mixed with a 50% contrast/saline solution (Omnipaque Iohexol, 350 mgI/mL, GE Healthcare), and infused intra-arterially in ∼0.1□mL pulses until luminal occlusion was confirmed by very slow flow of contrast at the occlusion site (total volume: 1-1.2□mL). The same procedure was adopted for the RM group with ∼200□µm SakuraBead® microspheres (CrannMed Ltd, Ireland), prepared by hydration in saline and contrast, then aspirated and injected under fluoroscopy until the same occlusion end-point was achieved as for the PM group. In both cases there was an observation period, followed by further injection of embolic as needed if flow persisted (total volume: 1.1-1.6□mL). The embolization was performed mostly in distal secondary branches where possible.

**Figure 1:**
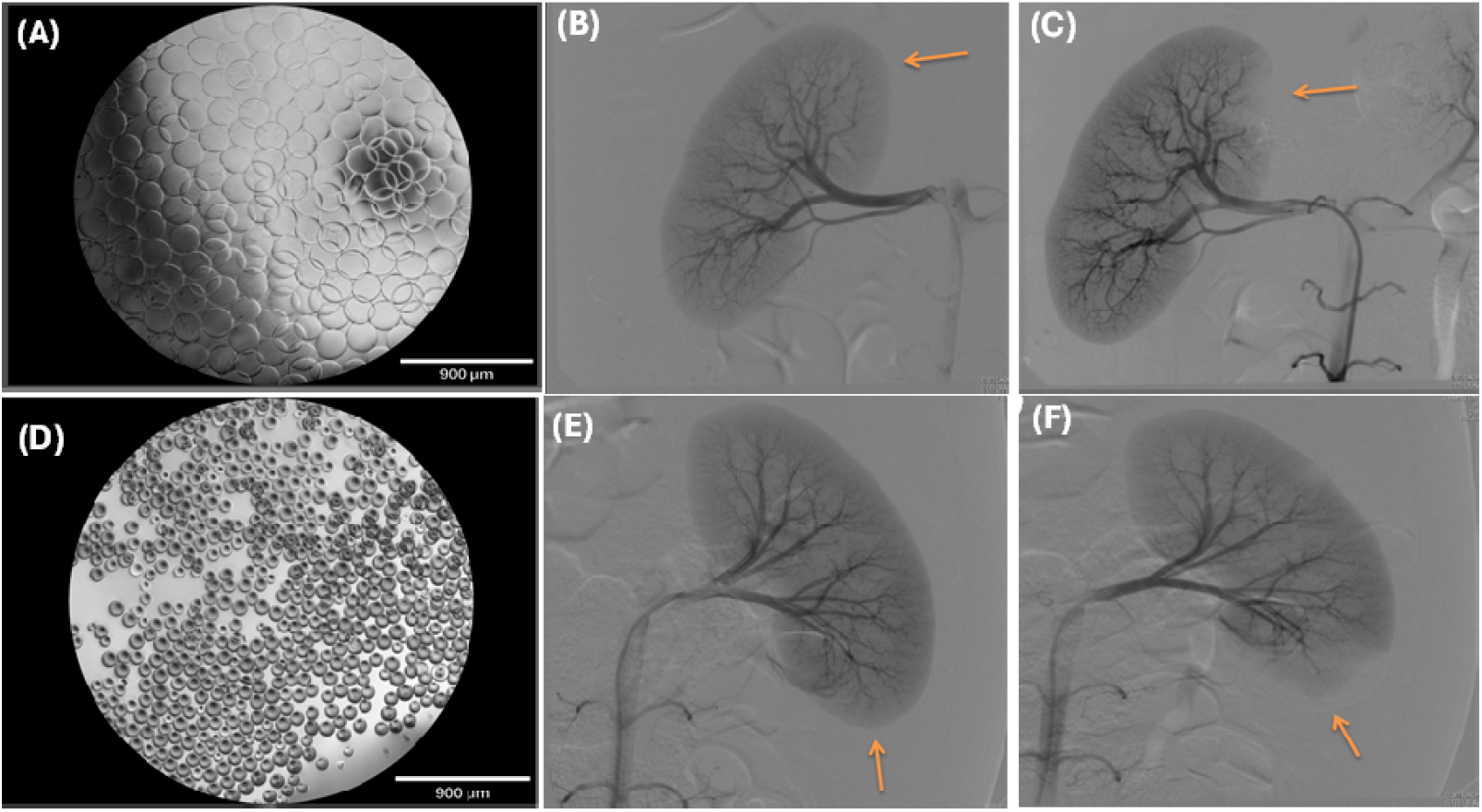
(A) Optical image of hydrated-RM; (B) Right cranial renal artery, before treatment with RM; (C) immediately post-treatment with RM, (D) optical image of PM; (E) Left caudal renal artery, before treatment with PM; (F) immediately post-treatment with PM. Arrows indicate embolization targets and resulting devascularization regions.

### 2.5 Histomorphometry

The kidneys were sampled at the proximal, middle, and distal treated region and at two sites of the untreated kidney region. Trimmed specimens were paraffin-embedded and stained with H&E to evaluate presence of embolic material, intrinsic vascular, local tissue and inflammatory responses. Ordinal histology was recorded by a qualified pathologist and data (scores) were plotted as mean ± SD. Statistical analysis was conducted for the RM group only and compared with the equivalent PM group for the same observation period. Two sampled t-tests were used for evaluating the significance difference (*p< 0.05, #p< 0.01and Δ p< 0.001). The macroscopic changes (for example, mild, minimal, absent etc.) that are used to explain the results are given in

## 3 RESULTS

### 3.1 Device Deployment and Clinical Observations

Figure 1(A) shows an optical micrograph of the RM hydrated in 1.5 ml saline for 2 minutes prior to use in a procedure, demonstrating complete hydration into smooth round microspheres of size 210 ± 8 μm (Fig 1A).

Representative angiographic images pre- and post-embolization for both RM and PM are shown in Figure 1 (B-C) and (E-F), respectively. Embolization of the desired targeted renal arteries was achieved in 100% of cases (as per Table 1) with a corresponding devascularized area of the distal parenchyma evident in all angiograms. All animals survived the procedure, maintained or gained weight until they were euthanized at pre-determined endpoints. For the evaluation of serum markers, only Creatinine Kinase showed elevated levels on Day 0 and 3 post-embolization as highlighted in Table 2.

**Table 2:** Representative data on serum markers and plasma coagulation parameters of animals treated with RM and PM evaluated at Day 0 (prior to intervention), Day1, Day 3, Day 7 and Day 28. (Bold value indication-out of the range).

| Embolitic beads | Follow up period | BUN mg/dL | CR mg/dL | GLUC mg/dL | NA mEq/L | K mEq/L | CL mEq/L | ALP U/L | ALT U/L | AST U/L | TBIL mg/dL | CK U/L | GGT U/L | TPRO g/dL | ALB g/dL | GLOB g/dL | CA mg/dL | PH mg/dL | CHOL mg/dL | TRI mg/dL | PT sec | APTT sec | FIB mg/ml |
| --- | --- | --- | --- | --- | --- | --- | --- | --- | --- | --- | --- | --- | --- | --- | --- | --- | --- | --- | --- | --- | --- | --- | --- |
| Reference range of parameters |  | 5-30 | 0.8-2.3 | 66-150 | 134-152.5 | 3.3-6.5 | 94-106.4 | 46-263 | 17-58 | 14-84 | 0- 0.5 | 100-416 | 10-60 | 5.2- 7.4 | 1.8- 4.0 | 2.6- 5.3 | 7.1-11.6 | 5.2- 9.6 | 28-95 | 16-39 | 12.8-16.9 | 18.1-27.2 | 100-400 |
| RM | DAY 0 | 9 | 1.1 | 73 | 139 | 3,7 | 100 | 134 | 33 | 17 | 0.1 | 212 | 30 | 5.2 | 2.9 | 2.3 | 10.2 | 9.1 | 55 | 18 | 13 | 13.2 | 95 |
|  | DAY 1 | 10 | 1.2 | 92 | 136 | 3.8 | 98 | 148 | 61 | 110 | 0.1 | **6640** | 36 | 5.4 | 3.1 | 2.3 | 9.4 | 7.4 | 61 | 35 | 12.5 | 13.5 | 191 |
|  | Day 3 | 8 | 1.4 | 109 | 143 | 3.8 | 103 | 123 | 29 | 13 | 0.1 | **606** | 26 | 5.9 | 3.7 | 2.2 | 10.5 | 8.5 | 79 | 31 | 11.4 | 12.6 | 172 |
|  | Day 7 | 12 | 1.2 | 94 | 140 | 4.3 | 100 | 149 | 23 | 13 | 0.1 | 249 | 34 | 5.4 | 3.0 | 2.4 | 249 | 34 | 5.4 | 30 | 11.2 | 12.2 | 160 |
|  | Day 28 | 14 | 1.6 | 100 | 140 | 4.1 | 102 | 123 | 35 | 24 | 0.1 | 392 | 36 | 5.6 | 3.1 | 2.5 | 9.5 | 7.6 | 79 | 29 | 11.6 | 12.0 | 109 |
| PM | DAY 0 | 8 | 0.9 | 88 | 135 | 3.7 | 100 | 92 | 30 | 21 | 0.1 | 339 | 21 | 5.3 | 3.0 | 2.3 | 9.8 | 7.8 | 54 | 27 | 14.1 | 14.2 | 78 |
|  | DAY 1 | 13 | 1. 2 | 93 | 136 | 3.8 | 102 | 129 | 24 | 16 | 0.1 | **8885** | 21 | 5.6 | 3.3 | 2.3 | 9.4 | 7.2 | 68 | 21 | 12.8 | 13.9 | 166 |
|  | Day 3 | 6 | 1. 3 | 100 | 144 | 3.9 | 102 | 129 | 24 | 16 | 0.1 | **567** | 24 | 6.2 | 3.8 | 2.4 | 10.5 | 8.3 | 106 | 19 | 11.8 | 11.5 | 104 |
|  | Day 7 | 6 | 1 | 79 | 137 | 4 | 100 | 103 | 29 | 18 | 0.1 | 228 | 30 | 5.4 | 2.5 | 2.9 | 9.0 | 8.8 | 58 | 29 | 12.6 | 12.9 | 136 |
|  | Day 28 | 11 | 1.4 | 94 | 138 | 4.0 | 101 | 132 | 32 | 15 | 0.1 | 396 | 28 | 5.4 | 2.9 | 2.5 | 9.2 | 7.1 | 97 | 27 | 11.5 | 10.3 | 137 |

### 3.2 Gross Necropsy Observations

In necropsy, treatment-related observations were only kidney specific. These were absent to mild in the RM group (Fig. 2 (A-D)), whereas extensive and consistent bilateral renal infraction was notable in the PM group across all time points (Fig. 2 (E-H)).

**Figure 2:**
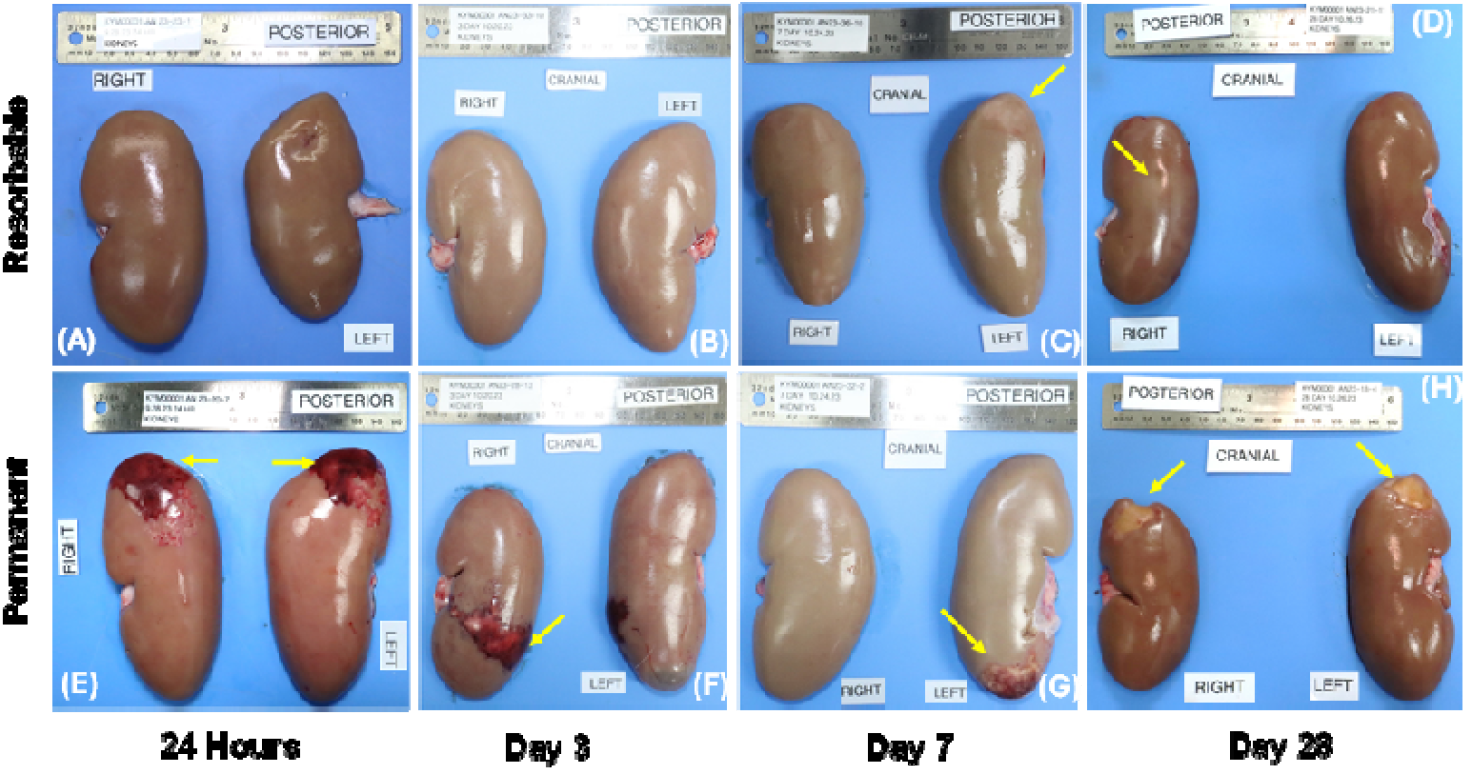
Gross kidney pathology post-necropsy. Top row (RM group): (A) 24h – no gross lesions; (B) Day 3 – pale focus (3×1□cm) on left kidney cranial pole, correlating with mixed histologic damage; (C) Day 7 – bilateral discoloration (2×1□cm left, 2×0.5□cm right); (D) Day 28 – no gross findings. Bottom row (PM group): (E) 24h – bilateral dark discoloration (cranial poles), indicating acute infarction; (F) Day 3 – red/purple areas near caudal poles (up to 5×2.5□cm), due to arterial occlusion; (G) Day 7 – dark caudal discoloration, chronic infarction; (H) Day 28 – sunken tan/brown areas (2×3□cm), indicating chronic infarction.

### 3.3 Histomorphometry: Intrinsic vascular response

In the RM group, no embolic material was observed beyond 24 hours, renal cortical necrosis was limited to 24 hours and Day 3, with significantly lower severity than in the PM equivalent group (p <0.01), as given in Figure 3(A). On the contrary, necrosis was consistently present across all the time points in the PM group. Fibrosis and fibroplasia appeared in the RM group from Day 3 onward but remained significantly lower than in the PM group on Day 28 (p□<□0.01) and from Day 3 onwards (p□<□0.01 and p□<□0.001). In the RM-treated group, no luminal occlusion was observed, and thrombosis occurred only on Day 3, with significantly lower severity than in the PM group equivalent (p□<□0.001).

**Figure 3:**
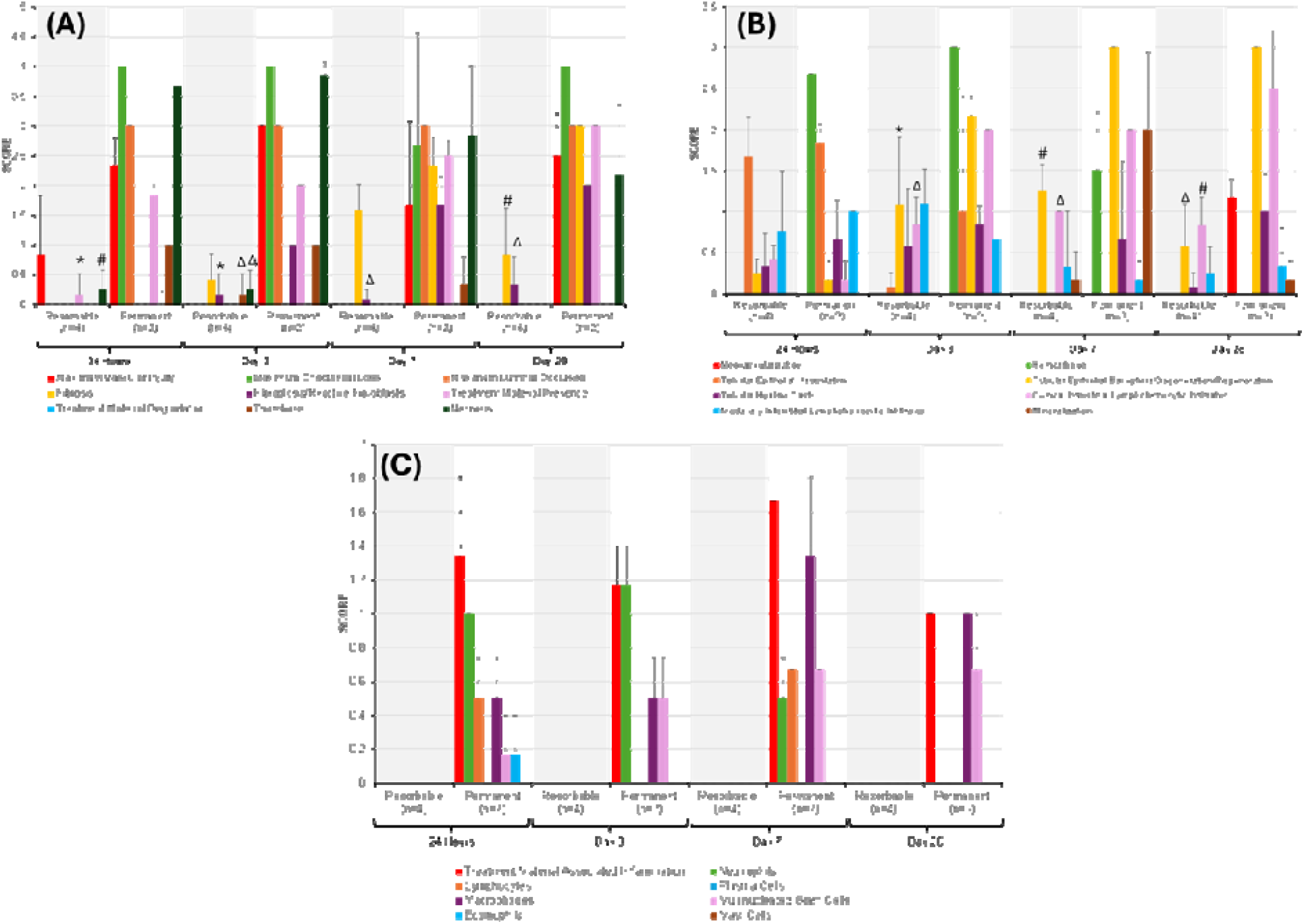
Histomorphology observations of RM and PM treated groups at Day 3, 7 and 28. (A) intrinsic vascular, (B) renal epithelial related and local tissue and (C) matory responses. (Statistical analysis: mean ± S.D, t-test significance *p< 0.05, #p< 0.01and Δ p< 0.00

### 3.4 Histomorphometry: Local Tissue Responses

From Figure 3(B), haemorrhage was not seen in the RM group, while in the PM group it resolved completely only on Day 28. In the RM group, tubular epithelial vacuolation resolved on Day 7, while tubular epithelial basophilia, vacuolation, hyaline casts, medullary and cortical interstitial lymph histiocytic infiltrates were present throughout the study period. Medullary and cortical lymph histiocytic infiltrates which are signs of chronic inflammation were significantly (*p <0.05, #p <0.01and Δp< 0.001) lower for RM than the corresponding PM group on Days 3, 7 and 28.

### 3.5 Histomorphometry: Local Inflammatory Response

There was no inflammation noted in untreated kidney sections as well as in the RM group, regardless of time point as shown Figure 3(C). Treatment material-associated inflammation in the PM group was persistent with neutrophils predominant through Day 3 and macrophages and giant cells at Day 7 and 28.

## 4 Discussion

### 4.1 Localized response to RM and PM embolization

Previous work has shown complete RM vascular recanalization of this product reported at 1-3hrs after use and compete degradation of breakdown products from adjacent tissue in the hours via gradual proximal refilling, indicating progressive microsphere breakdown [11]. This aligns with current findings, where no luminal occlusion was seen at 24 hours. In contrast, PM remained intact at all time points, consistent with its permanent nature [16]. By Day 3, RM-associated necrosis had resolved with tissue regeneration, while the PM group continued to show necrosis, vascular injury, and thrombus. RM induced a milder vascular response than PM. Compared to the PM treated group, the RM group induced milder localized ischemic stress and inflammation, with less tubular damage, fewer hyaline casts and vacuolation, and significantly lower levels of lymphocyte and histiocyte infiltration, tubular epithelial basophilia as along with complete absence of material-associated inflammatory response (Figure 3(B) and(C)). These findings highlight that RM’s ability to reduce localized ischemic and inflammatory, while enabling natural tissue recovery. This could be attributed to the rapid degradation of RM as well as the use of low immunogenic ultra-pure alginate used in the preparation of the RM [7]. It has been observed that the degree of embolic material present in lumen is related to the vascular inflammation and localized injury. Weng et al., showed that localized vascular inflammation persists until the microspheres degraded completely. [17][18]. This is consistent with the observations in this study, where a mild localized tissue reaction to RM is due to the rapid intravascular degradation of the beads, leading to reduced ischemia insult of the parenchyma around the occlusion site, whereas PM exhibited consistent luminal presence along with significantly higher vascular injury and inflammation.

### 4.2 Systemic response to RM and PM embolization

Upon embolization with RM or PM, most serum markers and all the haematology parameters remained within normal ranges, with only transiently early rise in creatine kinase which is likely due to mild and reversible ischemic injury. By day 3, CK levels declines as the recovery of the surrounding tissues progressed. Importantly, overall renal function (ALT, AST, BUN etc) remained unaffected, and both RM- and PM-treated animals-maintained blood homeostasis. These findings suggested that the breakdown products of RM unaffected the systemic parameters which are desirable properties for a resorbable embolic agent. Previously, we have estimated that breakdown products of RM last in human body for not more than 24 hours which could be attributed to the systemic-inertness of the RM[11].

### 4.3 Potential suitability of RM for MSK pain relief

Painful MSK condition stems from recurrent mechanical damage of joint tissue,causing abnormal neovessel and neoinnervation [1] [19]. This pathological loop can be disrupted by targeting neovascularity [20][21]. Neovessels are structurally abnormal and susceptible to ischemic damage [22]. Transient embolization using dissolvable imipenem/cilastatin particles has shown to necrotize neovessels effectively [23]. Given the transient nature and the induction of necrosis to normal renal arteries, of RM-may be a safe option for targeting neovessels in MSK disorder.

## 6 Conclusion

Embolization with SakuraBead RM appears safe, showing no signs of adverse tissue responses that would raise concerns for its use as a temporary occlusion agent. Tissue injury is mild and with no adverse sign of local or systemic inflammatory response in the porcine renal embolization model. These findings of the RM are aligned with the requirements of a temporary embolic agent suitable for use to treat pain associated with MSK pain conditions.

## ACKNOWLEDGEMENTS

This study was co-funded by the European Union and Enterprise-Ireland DTIF grant. Views and opinions expressed are however those of the authors only and do not necessarily reflect those of the European Union or European Innovation Council and SMEs Executive Agency (EISMEA). Neither the European Union nor EISMEA can be held responsible for them.

## Conflict of Interest statement

LF reports financial support was provided by Disruptive Technologies Innovation Fund (Grant number: 174147), Enterprises Ireland. LF reports a relationship with CrannMed Ltd that includes: board membership and employment. SA, PM and AL are inventors on the granted patent US 12083242 issued to CrannMed Ltd.

The IACUC Project Number for this study is I00362 (Global Protocol) ll.

